# A Convolution Based Computational Approach Towards DNA N6-methyladenine Site Identification and Motif Extraction in Rice Genome

**DOI:** 10.1101/2020.07.08.194308

**Authors:** Chowdhury Rafeed Rahman, Ruhul Amin, Swakkhar Shatabda, Md. Sadrul Islam Toaha

## Abstract

DNA N6-methylation (6mA) in Adenine nucleotide is a post replication modification responsible for many biological functions. Automated and accurate computational methods can help to identify 6mA sites in long genomes saving significant time and money. Our study develops a convolutional neural network (CNN) based tool i6mA-CNN capable of identifying 6mA sites in the rice genome. Our model coordinates among multiple types of features such as PseAAC (Pseudo Amino Acid Composition) inspired customized feature vector, multiple one hot representations and dinucleotide physicochemical properties. It achieves auROC (area under Receiver Operating Characteristic curve) score of 0.98 with an overall accuracy of 93.97% using 5 fold cross validation on benchmark dataset. Finally, we evaluate our model on three other plant genome 6mA site identification test datasets. Results suggest that our proposed tool is able to generalize its ability of 6mA site identification on plant genomes irrespective of plant species. An algorithm for potential motif extraction and a feature importance analysis procedure are two by products of this research. Web tool for this research can be found at: https://cutt.ly/dgp3QTR.

## Introduction

Different types of epigenetic modifications such as N4-methylcytosine, N6-methyladenine and 5-methylcytosine exist in DNA of different species^1^. DNA *N*^6^-methylation is one of them where methyl groups are transferred to DNA molecules^2^. It is one kind of post replication modification. Such modification has been identified in three kingdoms of life such as Bacteria, Archaea, and Eukaryotes^3^. *N*^6^-methylation occurs when methyl group is transferred to the 6^*th*^ position of purine ring of Adenine nucleotide. Such modification is responsible for discrimination between original DNA strand and newly synthesized DNA strand, gene transcription regulation, replication and repair^4,5^.

Laboratory based experimental methods such as sequencing (SMRT-seq, MeDIP-seq)^6^, methylated DNA Immunoprecipitation^7^, capillary electrophoresis and laser-induced fluorescence^8^ have been proposed to identify *N*^6^-methyladenine (6mA) sites in the genome. 6mA profile of rice has been obtained recently using mass spectrometry analysis and 6mA Immunoprecipitation followed by sequencing (IP-seq)^9^. Genome wide identification of 6mA sites is labor intensive and costly using such experimental methods. Computational experiment based pLogo plot^10^ on benchmark dataset regarding rice 6mA site identification shows nucleotide distribution to be different near 6mA and non-6mA sites^11^. So, accurate computational methods can be effective towards 6mA site identification.

i6mA-Pred was the first tool developed for 6mA site identification in rice genome using LibSVM package 3.18^12^ producing a high quality rice 6mA benchmark dataset containing 880 samples in each class. They used custom feature vector for each sample inspired by PseAAC (Pseudo Amino Acid Composition)^13^. 6mA-RicePred tool was developed for the same task utilizing the same dataset^14^. The development was based on SVM classifier utilizing Binary encoding, K-mer and Markov features. This same dataset was used to train and build iDNA6mA tool which utilized one hot matrix as sample feature^15^. The research used sequential one dimensional (1D) convolutional neural network (CNN) architecture for classification. Similar approach was used to develop SNNRice6mA tool^16^. The only difference was the use of group normalization layer in between convolution layers and an adaptive learning rate. The same benchmark rice genome dataset was used to build i6mA-DNCP tool^17^ utilizing di-nucleotide composition frequency and dinucleotide based DNA properties in a TreeBagging algorithm based classifier. Heuristic based methods were used for selecting four dinucleotide based DNA properties for classification. Same benchmark dataset was used for building SDM6A tool^18^ which utilized a two layer ensemble approach based on SVM and extremely randomized tree based models.

iDNA6mA-Rice tool was developed using a rice genome dataset containing 1,54,000 samples per class^11^ using one hot vector based linear feature space with Random Forest algorithm. MM-6mAPred tool utilized neighbour dependency information to detect 6mA site in rice genome leveraging Markov Model^19^. 6mA-Finder tool on the other hand was developed using the same algorithm utilizing seven sequence oriented information and three physicochemical property based features^20^. An optimal feature group was derived using recursive feature elimination strategy. Recently developed tools in this domain include DNA6mA-MINT tool^21^ that uses a hybrid of CNN and LSTM utilizing simple one hot encoding as feature. Another tool SpineNet-6mA^22^ used a hybrid of CNN and SpinalNet for rice 6mA site identification utilizing similar sequence encoding scheme. A recent study has evaluated and compared 12 publicly available N4-methylcytosine (4mC) site prediction tools on a large independent test set^23^. i4mC-ROSE^24^ and i4mC-Mouse^25^ are two of these tools. Tool i4mC-ROSE was developed for 4mC identification in Fragaria vesca and Rosa chinensis utilizing a random forest classifier using six encoding methods for feature extraction. Another tool i4mC-Mouse was developed for mouse genome 4mC site identification using six encoding schemes. They used six random forest based models for these six encoding schemes and then combined their confidence values linearly to make the final prediction.

Though the above mentioned methods achieve good performance, they lack the ability to learn the coordination between multiple types of sample feature representations for the classification task. All of them require either single feature vector^11,12,14,17–20^ or single feature matrix^15,16^ for their proposed models. While using multiple features, such requirement enforces manual feature concatenation and optimization irrespective of the heterogeneity and intrinsic value of the features. Our proposed deep learning based multi branch model has the capability of learning both the coordination and optimization of multiple heterogeneous features from training data without manual intervention. Simple extension of our model makes it capable of finding potential motifs. Intermediate output of our trained model can be used to find feature relative importance. Such properties are unique to our proposed model.

Our proposed i6mA-CNN model is a 1D CNN based branched classifier which can identify 6mA sites in plant genome. We work with four different types of feature matrices used for DNA sequence representation - PseAAC inspired customized feature matrix, monomer one hot matrix, dimer one hot matrix and dimer physicochemical properties. We use correlation based dimer physicochemical property selection technique in order to remove unnecessary and/ or redundant property features. Our proposed model is able to coordinate between these heterogeneous features and learn distinguishing features between 6mA and non-6mA sites successfully. We also perform LSTM sub-model based extension of our proposed model using the basic principal of attention mechanism^26^ and find potential motifs for 6mA site identification in rice genome. We believe such motifs to have potential importance for further research in this domain. Experimental comparison on two benchmark datasets shows the effectiveness of i6mA-CNN tool. Success on independent test datasets shows the robustness of our proposed tool.

In summary, our contributions are as follows:

- We provide a CNN model based state-of-the-art tool i6mA-CNN for 6mA site identification in rice genome.
- We show that our tool is able to generalize on other plants as well.
- We provide an attention based algorithm for potential motif finding extendable to any genome related identification task.
- We perform detailed feature analysis as part of our current research.

## Methods

### Benchmark dataset construction

Gene Expression Omnibus (GEO) database maintained by National Center for Biotechnology Information (NCBI) has a total of 2,65,290 6mA site containing sequences each of 41 base pair (bp) length with 6mA site at the center^27^. CD-HIT program^28^ with 80% (following similar research^11^) similarity threshold has been implemented on these sequences in order to avoid redundancy bias which resulted in 1,54,000 6mA site containing sequences. These are our positive set of samples. The negative samples have been obtained from NCBI^29^ by following three rules: (i) 41 bp long sequences with Adenine nucleotide at the center have been selected, (ii) It has been experimentally verified that the centered Adenine is not methylated, (iii) 6mA generally occurs near GAGG motif^30^. Statistical analysis shows that 17% of the positive samples contain GAGG motif. Negative samples have been selected such that this ratio remains similar among negative class samples; otherwise learning based model would distinguish based on this characteristics. Thus a large number of negative samples have been curated. For creating a balanced dataset, 1,54,000 negative samples have been randomly picked to form the negative class of samples^11,16,22,31,32^.

### Sequence representation

A sample sequence of our dataset looks as follows:

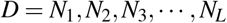

where, *L* is the length of the DNA sequence, which is 41 in our case and *N_i_* ∈ {*A, T,C, G*}. With appropriate mathematical representation of these sequences, deep learning models can learn class distinguishing features from the sequence local and global patterns^33^. We use the following sequence representations in our model:

#### PseAAC inspired feature representation

PseAAC (Pseudo Amino Acid Composition) has shown success in dealing with protein sequences^34,35^. Inspired by such success, PseKNC (Pseudo K-tuple Nucleotide Composition) has been introduced for dealing with DNA/RNA sequences in computational biology^36,37^. The general form of PseKNC has been designed in such a way that their values will depend on user specified feature extraction techniques from DNA samples. The four types of nucleotides of DNA can be classified into three categories of attributes. These attributes along with representation code have been provided in Table 1. The categories are as follows:

**Table 1.** Nucleotide categorical attribute-wise encoding

| Category | Attribute | Nucleotides | Code |
| --- | --- | --- | --- |
| Ring Structure | Purine | A, G | 1 |
|  | Pyrimidine | C, T | 0 |
| Functional Group | Amino | A, C | 1 |
|  | Keto | G, T | 0 |
| Hydrogen Bonding | Strong | C, G | 1 |
|  | Weak | A, T | 0 |

##### Ring number

A and G have two rings, whereas C and T have only one ring.

##### Chemical functionality

A and C are from amino group, whereas G and T are from keto group.

##### Hydrogen bonding

C and G are bonded to each other with 3 hydrogen bonds (strong), whereas A and T are bonded with only two (weak).

The above mentioned properties have significant impact on biological functions as well^38^. The *i^th^* nucleotide *N_i_* of a sequence can be represented by (*x_i_, y_i_, z_i_*), where *x_i_*, *y_i_* and *z_i_* represent ring structure, functional group and hydrogen bonding code (see Table 1) of nucleotide *N_i_*, respectively. Besides these three properties, we consider lingering density of each nucleotide to preserve sequence oriented features. Lingering density of nucleotide *N_i_* is defined below:

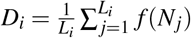

Here, *L_i_* is the length of the substring up to nucleotide *N_i_* and

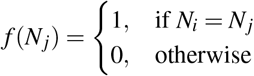

So, for each nucleotide, we have four properties to calculate. Since each sequence in our dataset is 41 bp long, we have a 41×4 dimensional feature matrix for each sequence. We illustrate this four property representation scheme on a toy example sequence “ACGGA” in Table 2. The resultant matrix representation is: [[1, 1, 0, 1], [0, 1, 1, 0.5], [1, 0, 1, 0.33], [1, 0, 1, 0.5], [1, 1, 0, 0.4]]

**Table 2.** Four property representation scheme detail of “ACGGA” sequence

| Pos | Nuc | X | Y | Z | D |
| --- | --- | --- | --- | --- | --- |
| 1 | A | 1 | 1 | 0 | $1/1 = 1$ |
| 2 | C | 0 | 1 | 1 | $1/2 = 0.5$ |
| 3 | G | 1 | 0 | 1 | $1/3 = 0.33$ |
| 4 | G | 1 | 0 | 1 | $2/4 = 0.5$ |
| 5 | A | 1 | 1 | 0 | $2/5 = 0.4$ |

#### Monomer one hot representation

Tool iDNA6mA was developed based on monomer one hot encoding^15^. 1D CNN has shown success in class discrimination based on one hot encoding. Each branch of our proposed model is also 1D CNN oriented. There are four types of nucleotides in DNA sequence and each of them is represented by a four size one hot vector. A, T, C and G are represented by (1,0,0,0), (0,1,0,0), (0,0,1,0) and (0,0,0,1), respectively. So, a 41 bp long sequence is represented by a 41 × 4 dimensional matrix.

#### Dimer one hot representation

There is statistically significant difference (considering p-value of 0.05) between 6mA and non-6mA site containing samples in terms of dinucleotide composition frequencies^17^. There are sixteen possible unique dimers such as AA, AT, AC, … in DNA sequences. We represent each dimer by a 16 size one hot vector. In a 41 length sequence, there are 40 overlapping dimers such as *N*_1_*N*_2_, *N*_2_*N*_3_, *N*_3_*N*_4_ …. So, each sample of our dataset is represented by a 40 × 16 dimensional matrix.

#### Dimer physicochemical property (DPP) based representation

Adenine methylation may have impact on DNA structure and its stability^39^. We have used the 90 DPPs used by PseKNC tool^36^. So, each dimer of our sample sequence is represented by a 90 size Z-score normalized vector, while each 41 bp long sample has been represented by a large 40×90 size matrix. If the DPP set contains features which include redundant or less important information, it may lead to overfitting^40^. The steps of DPP selection are as follows:

- We use Logistic regression model for 6mA site identification using one DPP at a time.
- We note the 5 fold cross validation Mathew’s correlation coefficient (MCC) score for each DPP and select the the top 20 DPPs. The higher the MCC, the better is the DPP for our identification task. MCC score is better than F1 score and accuracy for binary classification evaluation^41^. MCC score for top 5 DPPs are provided in Table 3.
- We select the topmost DPP. Now we want to remove redundant information. Each DPP is represented by a 16 size vector as there are 16 possible dimers. DPPs which have a high positive or negative correlation coefficient represent the same type of information. We calculate Pearson’s correlation coefficient for each of the 19 other selected DPPs with the topmost DPP.
- We leave out those DPPs which have correlation score greater than 0.9 (positive correlation) or less than −0.9 (negative correlation) with the topmost DPP.

**Table 3.** Five fold cross validation MCC score for the top 5 DPPs

| Dimer physicochemical property | MCC Score |
| --- | --- |
| Roll Rise | 0.58 |
| Major Groove Distance | 0.56 |
| Twist Stiffness | 0.53 |
| Protein Induced Deformability | 0.51 |
| Mobility to Bend Towards Major Groove | 0.49 |

Thus we select 9 DPPs out of 90 which include *Roll rise, major Groove distance, twist stiffness, protein induced deformability, mobility to bend towards major groove, melting temperature, Hartman trans free energy, protein DNA twist and tip.* We have also performed experiments using 18, 27, 36, 45, 54, 63, 72, 81 and 90 DPPs by including them in our proposed model. We have not obtained any performance gain in spite of the drastic increase of parameter number in our model.

### Model architecture

Our model architecture has been shown in Figure 1. There are four branches for four different feature matrices. Each branch has three 1D convolution layers put sequentially. The kernel size and filter number of each convolution layer have been shown in the figure. 1D CNN plays important role in position independent local feature extraction^42^. Each branch of the architecture learns class distinguishing features regarding its corresponding input feature matrix independent of the other three branches. The monomer one hot representation of the leftmost branch represents raw representation of sample sequence. Model parameters of this branch learn from this raw representation free from any feature specific knowledge bias. The second branch which has dimer one hot representation helps the model learn statistically significant differences between 6mA and non-6mA site containing samples in terms of dinucleotide pattern and frequency of occurrence^17^. The third branch uses DPP based representation. These properties play an important role in DNA specific classification and identification tasks^17,43,44^. This branch facilitates learning the interaction between these property values in a sequence order basis. The last branch is based on ring structure, functional group, hydrogen bonding and lingering density based nucleotide features. These features have impact on DNA biological functions^38,45^ enabling pattern learning from sequence coupled features.

**Figure 1.**
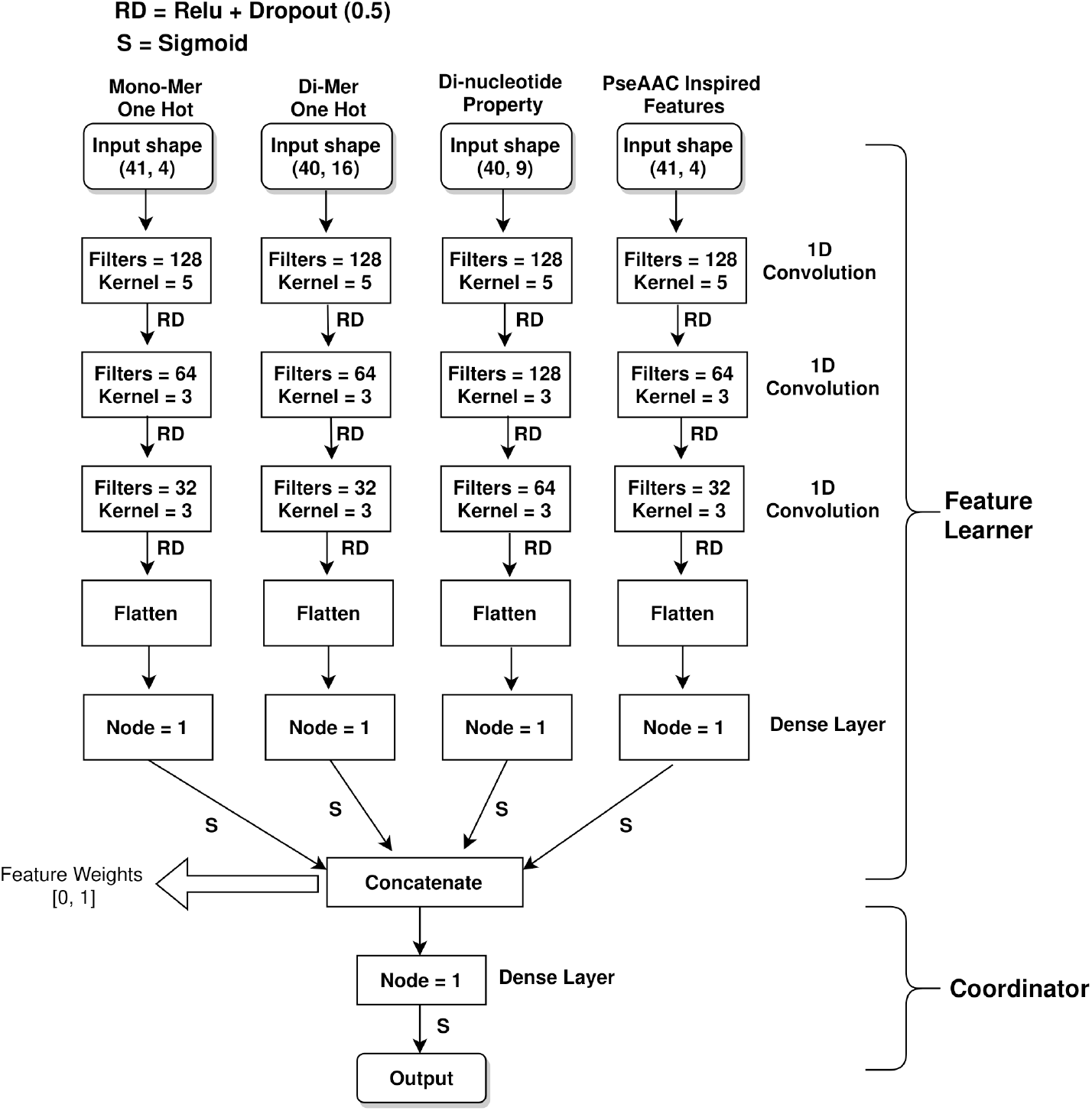
i6mA-CNN model architecture.

We need to have a way to combine these branches for a mutual classification decision. The CNN part of each branch i returns a matrix of dimension *m_i_ × n_i_*. We flatten and convert this matrix into a linear vector. We pass this vector through a dense node having Sigmoid activation function, which outputs a value in the range of 0 to 1. This single value obtained from the *i^th^* branch is its feature weight representative for 6mA identification task. Four feature weights from the four branches are concatenated and turned into a four size vector. This four size vector is the representative vector of all four branches. We denote this portion from input layer to feature weight concatenation layer as **Feature Learner**. The final dense layer with Sigmoid activation function provides the output value depending on the four values of the concatenated vector. If the output value is greater than 0.5, then we consider the sample sequence to have 6mA site. Else, we decide that there is no 6 mA site in the input sequence. This portion is our **Coordinator** of learned features.

### Performance evaluation

Accuracy (Acc), sensitivity (Sn), specificity (Sp) and Mathew’s correlation coefficient (MCC) have been used as evaluation metrics which are described as follows:

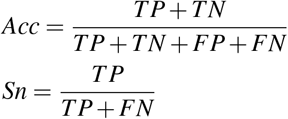

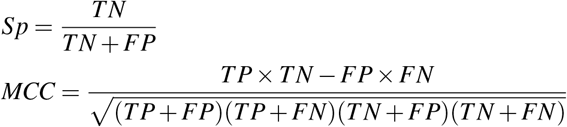

Here, TP, FP, TN and FN represent true positive, false positive, true negative and false negative, respectively. We consider the 6mA site containing samples as positive class samples. Apart from the above four metrics, we have also used area under receiver operating characteristic (auROC) curve for performance evaluation. This metric provides classifier performance irrespective of provided threshold for class discrimination. Acc, Sn, Sp and auROC lie in the range [0, 1], while the range for MCC score is [-1, 1]. Higher value indicates better classification ability.

### Potential motif finding

Specific local patterns frequently found in the samples of a class of nucleotide sequences are often important for identifying that particular class. Deep learning based classification models actively search for such potential motifs. We provide an algorithm for computationally identifying such potential motifs.

#### Attention model

We have constructed an attention based model shown in Figure 2 extending our proposed CNN architecture (shown in Figure 1). There are mainly two parts of our architecture - **Feature Learner** portion of our proposed CNN model and **bidirectional LSTM** based sub-model. The input for each time stamp of the LSTM sub-model is monomer one hot representation of a nucleotide of input sequence. So, there are total 41 time stamps in total. The LSTM portion gives us an output matrix **O** of dimension (41, 256) while the CNN gives us an output vector **C** of 256 size from **Dense Layer**. The dot product of **O** and **C** gives us a 41 length vector which represents the impact of each of the 41 nucleotides of the input sequence for 6mA identification. When passed through **Softmax** layer, these impacts normalize to a probability vector **S** of 41 length. This vector will represent the activation of each of the input nucleotides. When we take the dot product of **O** and **S**, we get a vector of 256 length. Mathematically, this dot product represents the weighted average of the 256 size output vectors over 41 time stamps of the LSTM, where the weight for the *i^th^* output vector is the *i^th^* index value of vector **S**. Thus the *i^th^* value of vector **S** represents the importance of the *i^th^* nucleotide input of the 41 length input sequence. Attention mechanism is about giving more importance to particular time stamp inputs than others. Finally, we concatenate the output vectors from both sides and pass the concatenated vector through a dense layer with Sigmoid activation function for 6mA identification as shown in figure.

**Figure 2.**
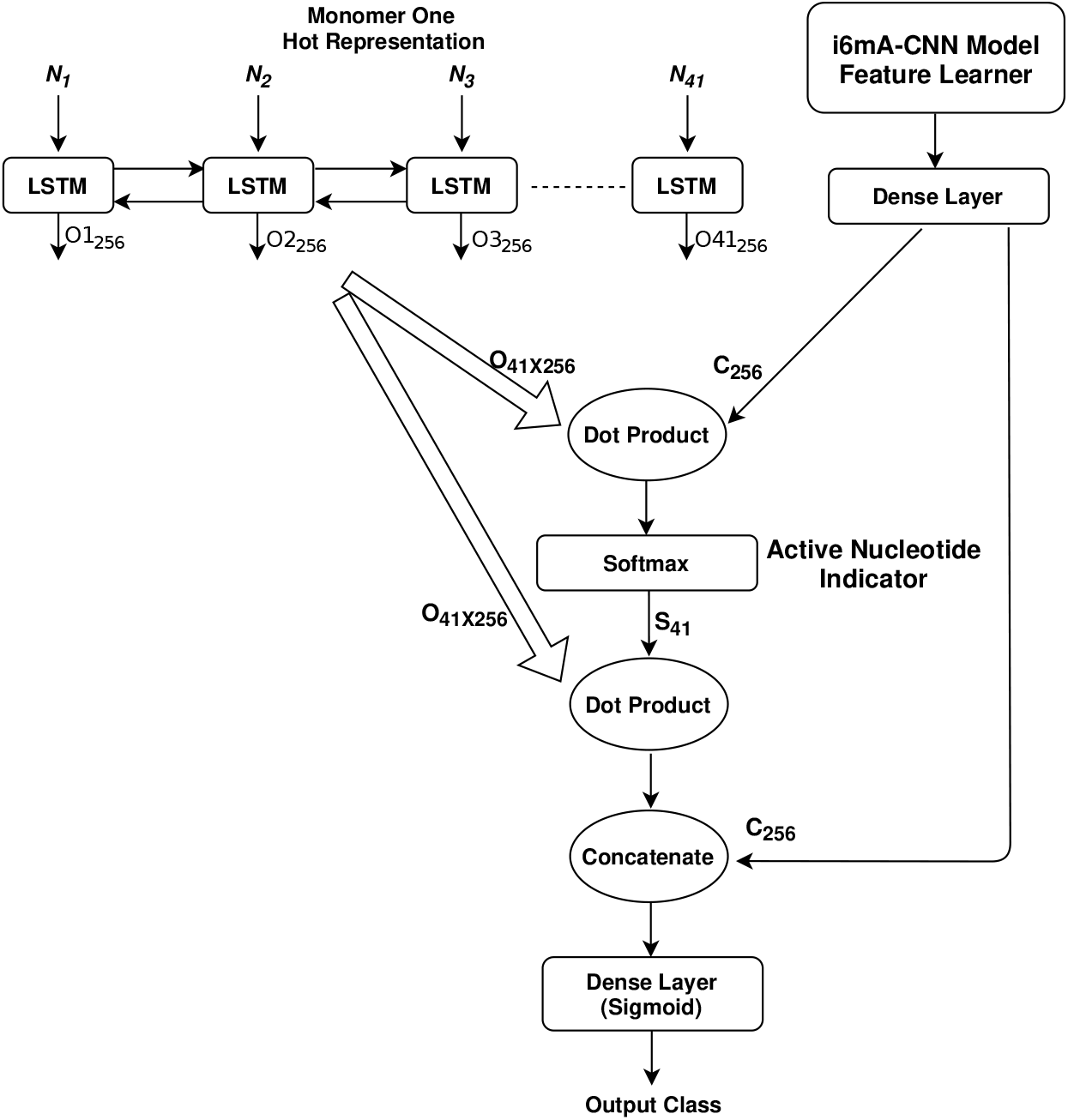
Attention based model for motif finding.

#### Training and utilizing attention model

We train our attention model end to end using our benchmark dataset. As our proposed attention model is based on **Feature Learner** portion of our proposed CNN architecture, this model provides similar performance on our benchmark dataset. The addition of LSTM based attention provides us with a 41 size vector **S** as intermediate output from **Softmax** layer as shown in Figure 2. When a 6mA site containing sequence is passed to our trained attention model, the *i^th^* index of **S** vector represents the probability of nucleotide *N_i_* of our input sample sequence to be activated for 6mA identification for this sequence. We utilize this intermediate output **S** for potential motif finding during 6mA identification.

#### Potential motif identification algorithm

When provided with a positive sequence as input, trained attention model provides a 41 size intermediate output vector **S** denoting a probability distribution. Each index probability value denotes the activation value of the corresponding index nucleotide of the input sequence. Any nucleotide which has an activation probability value greater than 1/41 is active during identification. In a uniform probability distribution, each nucleotide should receive 1/41 probability value in case it is neutral during classification. Consecutive positioning of active nucleotides denote an active region. When a particular nucleotide sub sequence is found to be active while identifying a large percentage of 6mA samples, that sub sequence can be labeled as a potential motif.

An illustrative example has been provided in Figure 3. There are three sequences in the example each of length 10. When we pass one of these sequences through our trained attention model, we get a 10 size intermediate output vector **S** denoting a probability distribution. **S** values have been provided for all three sequences in the figure. These activation values add up to 1 for each sequence. Nucleotides having activation value greater than 1/10 = 0.1 are active during identification. Following this notion, we see from figure that *CATAT*, *ATATG* and *CCATAT* are the active regions for Seq1, Seq2 and Seq3, respectively. *ATAT* is the common active region among these three. So, *ATAT* is the potential motif for this example.

**Figure 3.**
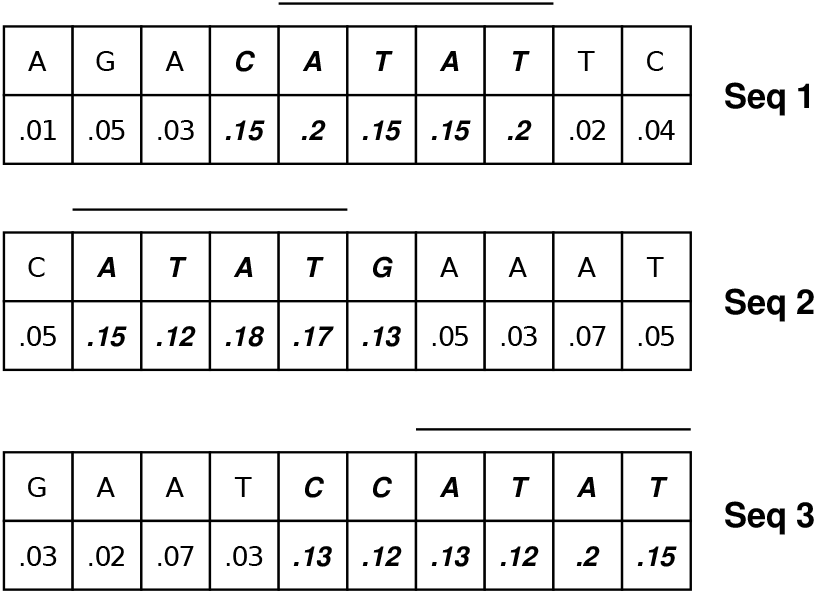
An illustrative example for motif identification.

#### Intuition behind algorithm correctness

Attention based model, when applied to sequence type data, learns which entities of the sequence are more important for reaching a conclusion or decision regarding output prediction from the sequence order information. Our designed model passes the weighted sum of the nucleotide monomers (monomer one hot vector) on to the final dense layer responsible for decision regarding class output. It is apparent that the nucleotides having higher weights are contributing at a larger scale towards the final decision. We choose active regions based on these weights assigned to the nucleotides by our model. These regions are indeed potential motifs used by the learning based model for deciding whether or not a sequence represents positive or negative class.

## Results

### Comparison with state-of-the-art tools on benchmark datasets

i6mA-Pred^12^, iDNA6mA-Rice^11^, iDNA6mA^15^, i6mA-DNCP^17^, MM-6mAPred^19^, SNNRice6mA^16^, 6mA-Finder^20^, 6mA-RicePred^14^, SDM6A^18^, DNA6mA-MINT^21^ and SpineNet-6mA^22^ are some of the state-of-the-art tools which can identify 6mA sites in rice genome. Consistency in dataset usage and evaluation method are necessary for comparison purpose with contemporary tools.

Benchmark dataset used in this research was used for constructing tool iDNA6mA-Rice, SNNRice6mA and SpineNet-6mA^11,16,22^ as well. Another smaller dataset containing 880 samples per class has been used by the other researches^12,14,15,17–21^ for benchmarking. They used either Jackknife testing or 10 fold cross validation for algorithm and model selection. We also validate our proposed tool on this dataset using similar validation methods. This smaller dataset has been obtained from NCBI GEO under accession number GSE103145. All sequences with 6mA site in the center and with modification score greater than 30 (according to Methylome Analysis Technical Note, 30 is the minimum score to call a nucleotide as modified) have been considered as positive samples. Each sample sequence is 41 bp long. CD-HIT^28^ with 60% similarity threshold has been applied on the chosen positive sequences in order to remove redundancy. Thus the 880 positive samples were obtained. The 880 negative samples of this dataset have been obtained in a manner similar to our negative benchmark dataset samples. As this dataset is small in size, it is not enough to train deep learning based models with large number of parameters. So, we have fine tuned our model on this small dataset training samples after training on our large benchmark dataset.

Detailed performance comparison between the tools used for 6mA site identification has been provided in Table 4. All the state-of-the-art tool results have been obtained from corresponding research articles^11,12,14–22^. Although the reported performance of the research article corresponding to *SpineNet-6mA* tool is superior to our proposed tool, through exact replication of the research we found out that the tool actually under performs (results provided in Table 4 for both datasets). The authors may have accidentally introduced some sort of training bias during SpineNet-6mA tool development. We have also performed Wilcoxon match pair test to examine if i6mA-CNN model shows significant performance improvement over SpineNet-6mA using the 10 MCC score pairs obtained from performing 10 fold cross validation on our benchmark dataset. We have performed a one tailed test as we want to see if our results are significant in a positive direction with a p-value of 0.01 and a null hypothesis that the median MCC score of both tools are identical. We obtain a p-value of .00256 which is less than 0.01. So, we reject the null hypothesis and reach the decision that i6mA-CNN provides significant improvement in terms of MCC score on our benchmark dataset. The same statistical test has been performed using Dataset 1 (contains 880 sample per class) of Table 4 using similar settings, where we get a p-value of 0.0002, which is far below 0.01. This improvement is even more significant.

**Table 4.** i6mA-CNN performance (marked in bold) comparison with state-of-the-art tools. **Dataset 1** is the 880 sample per class dataset used for comparison purpose, while **Dataset 2** is our benchmark dataset with 1,54,000 samples in each class. There are two results provided for SpineNet-6mA tool for both datasets - one obtained from reproducing the research and the other obtained from the reference research paper^22^

| Dataset | Evaluation Method | Tool | Sn | Sp | Acc | MCC | auROC |
| --- | --- | --- | --- | --- | --- | --- | --- |
| Dataset 1 | Jackknife Testing | i6mA-Pred | 83.41% | 83.64% | 83.52% | 0.67 | 0.91 |
|  |  | **i6mA-CNN** | **90.61%** | **94.12%** | **93.15%** | **0.85** | **0.98** |
| Dataset 1 | 10 Fold Cross Validation | iDNA6mA | 86.7% | 86.59% | 86.64% | 0.73 | 0.93 |
|  |  | i6mA-DNCP | 84.09% | 88.07% | 86.08% | 0.72 | 0.93 |
|  |  | MM-6mAPred | 89.32% | 90.11% | 89.72% | 0.79 | — |
|  |  | 6mA-Finder | — | — | — | — | 0.94 |
|  |  | SDM6A | 85.2% | 90.9% | 88.1% | 0.76 | 0.94 |
|  |  | 6mA-RicePred | 84.89% | 89.66% | 87.27% | 0.75 | — |
|  |  | DNA6mA-MINT | 94.25% | 90.8% | 92.53% | 0.85 | 0.95 |
|  |  | SpineNet-6mA (reproduced) | 86.48% | 92.39% | 89.43% | 0.79 | 0.95 |
|  |  | SpineNet-6mA (from reference) | 93.75% | 95.79% | 94.77% | 0.89 | 0.98 |
|  |  | **i6mA-CNN** | **90.35%** | **94.62%** | **92.48%** | **0.85** | **0.98** |
| Dataset 2 | 5 Fold Cross Validation | iDNA6mA-Rice | 93% | 90.5% | 91.7% | 0.83 | 0.96 |
|  |  | SNNRice6mA | 94.33% | 89.75% | 92.04% | 0.84 | 0.97 |
|  |  | SpineNet-6mA (reproduced) | 93.94% | 91.36% | 92.15% | 0.86 | 0.97 |
|  |  | SpineNet-6mA (from reference) | 95.71% | 92.92% | 94.31% | 0.88 | 0.98 |
|  |  | **i6mA-CNN** | **95.13%** | **92.81%** | **93.97%** | **0.88** | **0.98** |

Tool i6mA-CNN achieves a high average 5 fold cross validation auROC score of 0.98 on our benchmark validation set. ROC curves on all 5 fold validation sets have been shown in Figure 4. The red dotted straight line denotes ROC curve for models with auROC score of 0.5 which denotes random prediction performance. The 5 curves obtained from 5 folds of validation are all high above this red straight line denoting quality prediction. All 5 curves are overlapping which shows the consistency of our proposed model performance in all 5 folds.

**Figure 4.**
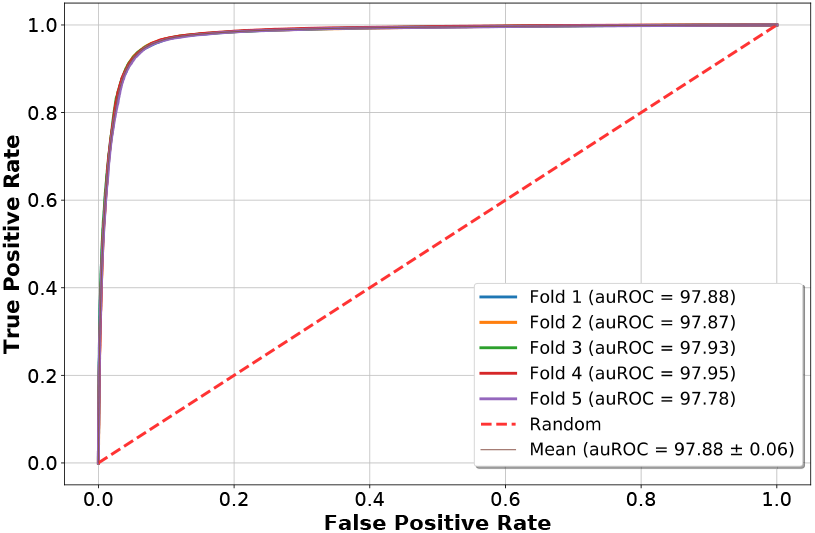
Five fold cross validation ROC curves on benchmark dataset.

### Performance on independent test sets

Independent test set performance of a tool can help in proving the generalization ability of that tool. Following procedures similar to our benchmark dataset construction, we have used three independent test datasets in order to evaluate the robustness of our proposed tool in terms of *N*^6^-Methyladenine site identification in plant genome. Fragaria vesca (wild strawberry) and Rosa chinensis (Chinese rose) 6mA site containing sequences have been obtained from the MDR database^46^, while Arabidopsis thaliana (cress weed plant) sequences have been obtained from NCBI GEO with accession number GSE81597^47^. Only the sequences having Adenine nucleotide at the center with modification score of at least 30 have been kept. Then CD-HIT software has been implemented for removing sequences having 80% or more similarity. Results provided in Table 5 show the potential of accurate 6mA site identification of our tool irrespective of plant species. This same dataset was used as independent test set for a recently developed tool i6mA-DNCP^17^. The tool achieved a success rate of 96.63%, 91.51% and 91.89% on Fragaria vesca, Arabidopsis thaliana and Rosa chinensis dataset, respectively. Although our tool outperforms i6mA-DNCP in terms of the success rate in the first two species, our tool under performs on Rosa chinensis dataset. The reason is that our tool is deep learning based and requires a lot of training samples. Since the sample number in Rosa chinensis 6mA dataset is only 1126, our model learning ability in this case has not been fully expressed.

**Table 5.** 6mA site identification performance of i6mA-CNN on three plant genomes

| Plant Genome | Number of Samples | Number of Correct Predictions | Success Rate |
| --- | --- | --- | --- |
| *Fragaria vesca* | 8979 | 8780 | 97.78% |
| *Arabidopsis thaliana* | 67650 | 64800 | 95.78% |
| *Rosa chinensis* | 1126 | 966 | 85.79% |

### Model validation on imbalanced training dataset

Our utilized benchmark dataset is balanced. But in real life, most genome classification datasets are highly imbalanced, where negative samples dominate the dataset. In such cases, learning based models using redundant features and large number of parameters tend to be biased towards the dominant negative class. While performing 5 fold cross validation on our benchmark dataset, we implement random undersampling on the positive training samples and show performance for 1:1 (balanced), 1:5, 1:10 and 1:20 (highly imbalanced) positive to negative training sample ratio in Table 6. Our model shows reasonable performance even when the imbalance ratio is 1:20. As the ratio of negative samples increases, the gap between Sn and Sp score increases, while MCC score decreases - a typical symptom of increase of bias towards dominant class samples. The gap between Sn and Sp score is only 18%, even when positive to negative training sample ratio is as high as 1:20. With the presence of 20 times more negative samples than positive samples, i6mA-CNN still manages to achieve an auROC score higher than 0.90.

**Table 6.** 5 fold cross validation performance of i6mA-CNN model when trained with disproportionate number of positive and negative samples

| Positive Negative Sample Ratio | Sn | Sp | Acc | MCC | auROC |
| --- | --- | --- | --- | --- | --- |
| 1:1 | 95.13% | 92.81% | 93.97% | 0.88 | 0.98 |
| 1:5 | 91.08% | 95.55% | 94.81% | 0.82 | 0.94 |
| 1:10 | 84.7% | 97.1% | 95.98% | 0.77 | 0.92 |
| 1:20 | 80.23% | 98.2% | 96.02% | 0.73 | 0.91 |

### Feature significance analysis

Our proposed CNN based architecture shown in Figure 1 uses four heterogeneous feature matrix representations. As shown in Figure 1, the **Feature Learner** portion of our proposed architecture passes a concatenated vector of size 4 on to the **Coordinator** portion, where each value of this vector is in range of 0 to 1. Let us name this vector as *Confidence_Vector*. When a sample sequence is passed to our trained model, *i^th^* index value of this *Confidence_Vector* represents the weight or confidence of *i^th^* input feature matrix among the 4 input matrices for 6mA identification in this sample. The 4 input matrices represent -(i) monomer one hot representation (Monomer), (ii) dimer one hot representation (Dimer), (iii) physicochemical property representation (PhyChe) and (iv) PseAAC inspired feature matrix (PseAAC). The Y-axis of Figure 5a denotes the percentage of positive samples in terms of frequency. The bar plot shows the percentage of positive samples to produce low, medium and high value in *Confidence_Vector* for the 4 heterogeneous features. For example, from Figure 5a it is clear that no positive sample produces low Monomer confidence score, very few of them produce medium range Monomer confidence score, while almost all positive samples produce high Monomer confidence score. So, Monomer plays a significantly more important role for positive sample identification, while PhyChe representation plays a minor role, because almost all samples produce low confidence score for this property. Similar plotting has been shown in Figure 5b for negative sample. The significance of the features are quite different for negative samples. Dimer and PseAAC receive high value for most samples, while Monomer is on the low side of significance in most cases.

**Figure 5.**
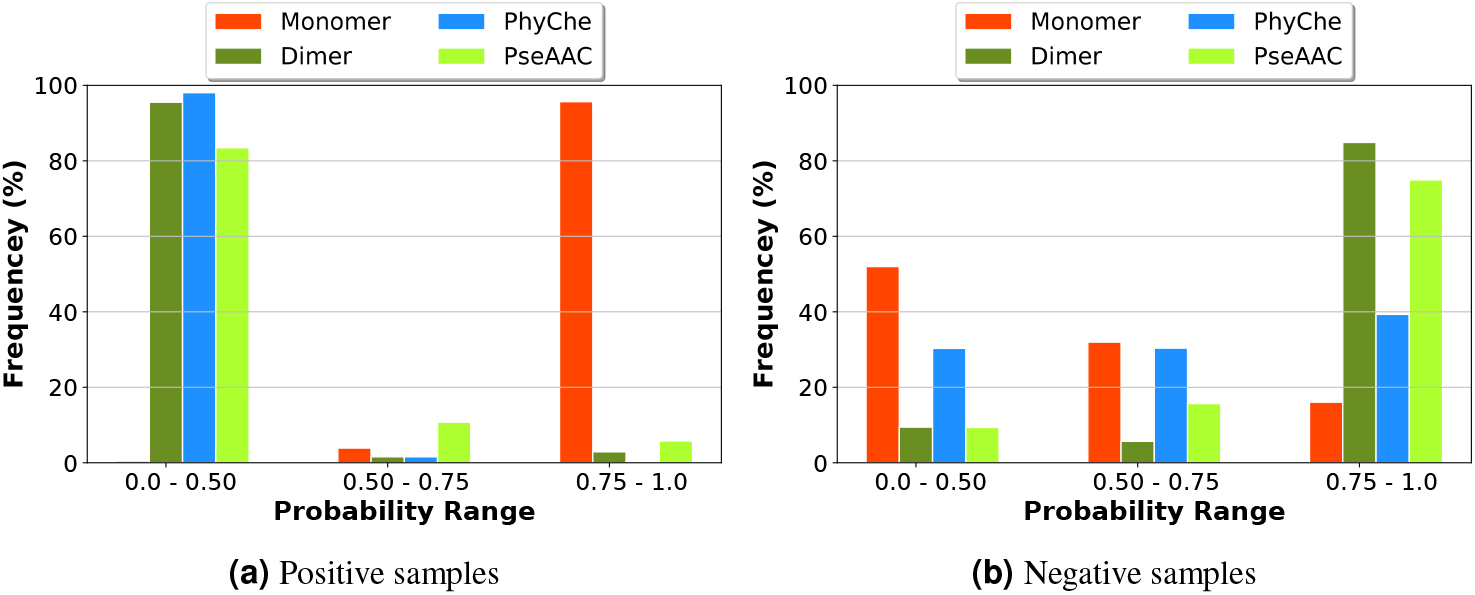
Feature significance analysis for 6mA site identification in positive and negative samples of benchmark dataset.

We have also performed experiment using weighted ensemble learning based approach, where we use a simple sequential convolutional neural network and use only one of the four features for with each network. Implementing weight based ensemble learning has provided us with five fold cross validation Sn, Sp, Acc, MCC and auROC score of 93.78%, 91.54%, 92.66%, 0.86 and 0.96, respectively on our benchmark dataset. The reason for this under performance compared to our coordination based proposed CNN model is that ensemble learning takes each feature separately and there is no scope of learning any mutual coordination between them from training data.

As part of ablation study, we have experimented with single feature using simple sequential 1D CNN based model instead of using all four of our features together. Performance comparison between each independent feature and all four features combined using i6mA-CNN model has been shown in Table 7. Experimental results indicate that there are a significant amount of samples whose classification depends on the coordination of multiple features together.

**Table 7.** Performance comparison when each individual feature is used independently. Experiments have been performed on the benchmark dataset containing 880 samples in each class.

| Features Used | Sn | Sp | Acc | MCC | auROC |
| --- | --- | --- | --- | --- | --- |
| Monomer | 87.27% | 85.8% | 86.53% | 0.73 | 0.94 |
| Dimer | 87.3% | 86.48% | 86.8% | 0.74 | 0.94 |
| PhyChe | 85.36% | 86.27% | 85.8% | 0.72 | 0.93 |
| PseAAC | 85.47% | 83.98% | 85.22% | 0.70 | 0.92 |
| All Four Features | **90.35%** | **94.62%** | **92.48%** | **0.85** | **0.98** |

### Identified potential motifs

Potential motifs are patterns which learning based algorithms actively search in order to perform identification tasks. A list of potential motifs found for 6mA site identification on our benchmark dataset have been provided in Table 8. The top motif that we have found is **ATAT**. It was found to be active 5.2% of the time during 6mA site identification. Experimental validation shows that 6mA occurs most frequently at GAGG motifs^30^. Our computational method also shows GAGG as the third most significant motif used by our learning based model to identify 6mA site. Future research can be conducted on experimental verification of rest of the potential motifs mentioned in Table 8.

**Table 8.** Potential motifs for *N*^6^-methyladenine site identification

| Motif Sequence | Active Occurrence % |
| --- | --- |
| ATAT | 5.2 |
| AGGC | 4.1 |
| GAGG | 4.0 |
| AAAA | 3.9 |
| AAAT | 3.5 |

### Web server implementation

We have used Tensorflow library, Numpy array, Pandas data processing tool, Sklearn library and Flask for i6mA-CNN tool development. Our developed web server is freely available at: https://cutt.ly/dgp3QTR. The web server requires sequences to be provided in FASTA format. Users can also type in sequences. Each sequence has to be 41 bp long. The predictions are made by our trained proposed CNN model in the backend. Our web server prediction time is fast because of specialized batch prediction by the model. The predictions can be easily downloaded in .csv format. A user can provide maximum 1000 sequences at a time for prediction in the user friendly web server. In case a user wants prediction for more than 1000 sequences at a time, a Google Colaboratory based server is also provided. Even for a million number of sequences, this Colab based server takes less than a minute to finish providing the prediction results. The Colab based server is freely available at: https://cutt.ly/hgdB09A.

## Discussion

We propose a tool i6mA-CNN aimed at automated 6mA site identification in rice genome. Such post replicational epigenetic modification plays a key role towards biological functions. We use a four branch convolutional model having the ability to coordinate among four heterogeneous feature matrices of a single genome as the backbone of our proposed computational tool. This model architecture is easily extendable to additional heterogeneous feature matrices. We also provide an attention based computational algorithm for potential motif identification which can be used to retrieve a probable list of motifs for any genome related identification task. We further show detailed feature analysis using intermediate output of our proposed model which can be used in any genome related classification or identification task.

Our approach achieves impressive results on two benchmark datasets for rice 6mA site identification. We use jackknife testing, 10 fold cross validation and 5 fold cross validation as means of performance evaluation on the benchmark datasets. We have achieved an impressive MCC score of at least 0.85 and an auROC score of approximately 0.98 in all our evaluation methods on both benchmark datasets. We have also tested our tool on three different plant genome 6mA site sequence datasets other than rice, where our success rate was over 95% for two of these independent test datasets. Even the worst success rate was over 85%.

The backbone model of i6mA-CNN tool is a CNN based branched model devoid of recurrent layers. Recurrent layers have not been used for several reasons. First, it is not possible to provide multiple feature representations simultaneously in each time stamp input in recurrent neural networks (RNN). Second, each time stamp of such networks only accept linear vector representation which would limit our current ability of using matrices. Third, RNNs give importance to positional information^48^. But classification of genome sequence often depends on the presence of potential motif type patterns irrespective of their position in the sequence^30^. Finally, training and validation using recurrent layer based networks are costly because of sequential type of computation. 1D CNN based model having multiple convolution layers have the capability of encoding and learning both short and long range patterns. Parallel processing is possible in such networks which allow fast computation. Because of such inherent characteristics, our proposed tool is fast, accurate and generalizes well to other plant genomes for the same task. Although 1D CNN model was used in iDNA6mA^15^, SNNRice6mA^16^, DNA6mA-MINT^21^ and SpineNet-6mA^22^ tool development, they had either used a simple sequential architecture or a CNN-LSTM hybrid having no capability of pattern coordination learning between multiple heterogeneous features.

One may argue that i6mA-CNN model is costly in terms of run time compared to simple learning based models such as SVM. With the emergence of GPU based parallel matrix computation, run time is no more a problem, especially when there is provision for batch-wise prediction. We have recorded the run time of both SVM^49^ and i6mA-CNN model for 6mA site identification in 1000 sample sequences each of 41 length. SVM and i6mA-CNN model take 0.1218 seconds and 0.1222 seconds, respectively for this bulk prediction task. Our model is only 0.3% slower than SVM in terms of prediction speed which is negligible in a setting where we do not want real time online prediction.

Proposed i6mA-CNN tool is expected to be an effective tool for automation in epigenetics related research. Future research should aim at developing an automatic feature extraction based model architecture capable of showing satisfactory performance for any epigenetics related identification task provided that it is given appropriate training sample sequences as input. Such a tool is expected to eliminate the hassle of manual feature selection and appropriate model selection for each separate epigenetics related identification task. Experimental verification of the identified potential motifs in this research is also a possible direction for future research.

## Availability of materials and data

- All preprocessed datasets used in this research including two benchmark datasets and three independent test datasets are freely available at: https://cutt.ly/Kgd0kUD.
- Documented code for i6mA-CNN tool model building, training and 5 fold cross validation on our benchmark dataset is available for use at: https://cutt.ly/Ygd0Jdt
- Code for attention model building, training and motif finding algorithm is available for use at: https://cutt.ly/zgjecnj.
- Supplementary material consists of di-nucleotide physicochemical property values and the model .h5 file (used as the prediction engine for i6mA-CNN tool). They have been provided for use at: https://cutt.ly/igd37Gi

## Author Contribution

C.R.R and S.S. conceived the research. C.R.R. designed the algorithm and the models. R.A. contributed to code implementation and experimentations. C.R.R. wrote the manuscript, while S.S. performed critical review. M.S.I.T. developed the web tool with the assistance of R.A. All authors reviewed and approved the manuscript.

## Notes

https://cutt.ly/Kgd0kUD

https://cutt.ly/Ygd0Jdt

https://cutt.ly/zgjecnj

https://cutt.ly/dgp3QTR

### Competing Interest Statement

The authors have declared no competing interest.

### Summary of Updates

Motif finding method added; results updated; discussion added; web server added; figures revised.

